# Deep Learning for Predicting Stem Cell Efficiency for use in Beta Cell Differentiation

**DOI:** 10.1101/2025.05.22.652867

**Authors:** Franziska J. Schöb, Alexander Binder, Valentina Zamarian, Valeria Sordi, Hanne Scholz, Anders Malthe-Sørenssen, Dag Kristian Dysthe

## Abstract

Recent clinical trial data have shown that cell therapy holds curative potential for type-1 diabetes, however the large amounts of lab-grown cells required is a substantial bottleneck. The necessary cell differentiation process exhibits substantial variability, even among clones of induced pluripotent stem cells generated from the same patient. In contrast to many established data sets in medical imaging, human experts struggle to see the difference between highly- and lowly-efficient cell clones. We therefore propose an image-based deep learning model to guide the selection of the most efficient stem cell clones.

We apply different deep learning model architectures to learn the morphological differences between good and bad stem cell clones and classify them based on only phase-contrast imaging. To gain critical insight into the learned features and enhance trust in our model, we use layer-wise relevance propagation (LRP), and Fourier-based frequency analysis.

Using an EfficientNet-V2-S model, we obtain a novel early prediction for the outcome of the differentiation process from patient-derived stem cell to β-cells using only phase-contrast images. Clone level accuracy is 76.7 *%* at 24 hours and 96.7 *%* at 53 hours after start of differentiation. The LRP-generated attribution maps and structure factor analysis show that the structure of the cell population is an important predictive feature.

All in all, we present a highly predictive model for successful stem cell differentiation from phase-contrast images, which learns biologically relevant features. This study is a proof-of-concept that deep learning combined with label-free imaging can guide selection of induced pluripotent stem cell clones, thereby reduce cost of β-cell production, and bring curative treatments one step closer to the clinic.

## 1. Introduction

1 in 10 adults were living with diabetes in 2022 globally with a rising trend.^1^ Diabetes mellitus type 1 (T1D) is a chronic autoimmune disorder, defined by loss of pancreatic β-cells leading to insulin deficiency and thereby loss of the ability to regulate blood glucose levels.^2^ Patients therefore require regular injections of insulin during their whole life.^3^

Stem cells are a cell type which has the ability to self-renew and differentiate into other specialized cell types, opening up the possibility to utilize them in regenerative medicine and cell therapies. One promising application of cell therapy is T1D, where transplantation of functional β-cells could restore normal insulin production.^4^ First clinical results have recently confirmed this potential^5^.

Ideally, β-cells should be produced from patient-derived induced pluripotent stem cells (iPSC) in order to avoid immune rejection after transplantation. Based on previous studies on pluripotency induction,^6,7^ procedures have been developed to differentiate iPSC into β-cells.^8^ This differentiation process takes about 34 days, is costly and labor-intensive. The yield of functional β-cells is highly variable both inter- and intra-experimentally.^9,10^ In addition, the cost of cell therapy is substantial, making variability in experimental outcome an even bigger challenge.

Today, deep learning is trending in the medical domain including medical imaging, however it is most often used for CT, MRI, and other clinical imaging modalities.^11,12^ More recently, we see it used for microscopic imaging of cells, often with the goal of segmentation and tracking.^13–15^

Here, we present a deep learning-based approach to predict cell fate based on microscopy images. Our model predicts successful experimental procedures already during the first four days of differentiation based on phase-contrast imaging. It involves a novel dataset of a single cell type (iPSC) changing over time, which is distinctively different from most other common datasets, such as tissue sections which contain many different cell types at one time. Our approach aims at acceleration of cell production in vitro by providing a criterion when to restart the process with another batch. In contrast to traditional methods for cell fate analysis, such as gene expression measurements or staining for specific markers, our live-cell imaging approach is non-invasive, meaning it does not require fixation or staining of the cells. This is especially important in this case where available cell material is very limited. Also, it allows us to monitor the same cells over extended time periods without disruptions. We focus on the first four days of the 34 day differentiation, which should result in Definitive Endoderm (DE) cells, also known as stage 1.^16^

The task of predicting cell fate has been demonstrated as possible before on other cell types,^17–19^ however most of these studies focus on classifying between differentiating and non-differentiating cells and differentiation towards separate cell types, or rely on different image modalities like fluorescence-stained or Raman spectroscopy images. Here, we classify only differentiating cells towards the same fate into a low and a high expression group. Morphological changes during differentiation are clearly visible at much later times, but differences between clones at such early time points are difficult to see even for a trained expert. Selection of the best cell clones for further differentiation and use in cell therapy is therefore challenging and requires new approaches like ours for support.

## 2. Methods

All data and code will be available at zenodo and github, respectively.

### 2.1. Data preparation

Cell culture: iPSC differentiation Six patient-derived iPSC clones were differentiated to Definitive endoderm (DE) stage using the STEMdiff Definitive Endoderm Kit (STEMCELL) following manufacturers instructions. The differentiation was carried out as biological duplicates in 6-well plates.

Image acquisition of cell cultures We utilized the IncuCyte^®^ Live-Cell Analysis Systems (Sartorius), an automated microscope operating inside an incubator, ensuring optimal cell culture conditions and continuous imaging. 121 phase-contrast images per well were acquired for the first four days of differentiation (until DE stage) at 1 hour intervals. 136,488 1408 × 1040 pixels, 8-bit gray-scale images were obtained in total.

Ground truth label generation via flow cytometry For flow cytometric analysis, DE stage cells were trypsinized with 1x Trypsin (Lonza) for 5 min at 37 °C and stained for CXCR4, a DE stage specific marker. The cells were stained with a live/dead Fixable Near-IR Dead Cell Stain Kit (Life Technology). For CXCR4 staining, cells were incubated with antibody for 15 min at RT, then washed with FACS buffer and fixed with Cytofix/Cytoperm (BD Biosciences) for 20 min at 4 °C. Samples were then run through a FACS Canto Cytometer (BD) and cell counts were analyzed with FlowJo 10.8 (BD Life Sciences). Measured CXCR4 expression levels are shown in Table 1.

| Clone | 103 | 104 | 109 | 110 | 116 | 117 |
| --- | --- | --- | --- | --- | --- | --- |
| CXCR4 | 92.2 | 89.2 | 68.6 | 8.0 | 96.5 | 73.0 |

Dataset splitting Our dataset consists of the above described phase-contrast images and flow cytometry measurements of CXCR4 expression as ground truth labels. High levels of CXCR4 expression represent cells in the Definitive Endoderm stage, which is a measure of successful differentiation.

We decided to go for a classification setting since we had only 6 independent CXCR4 measurements. We defined two classes based on a CXCR4 expression with a threshold of 84.0% based on through biological analysis of the 6 clones. This analysis revealed that there are two distinct classes of clones, i.e. bad and good ones. The two classes have been shown to have differential expression of genes involved in stemness and trilineage specification as well as distinct success rates of differentiation into β-cells. These biological results will be available in a manuscript which is in preparation. The percentage of CXCR4-expressing cells correlates with the efficiency of final differentiation into functional mature β-cells; therefore, CXCR4 has been used in this study as a proxy for all differentially expressed genes. Clones with an expression higher than 84.0% represent class 1, while a lower CXCR4 expression represents class 0. Our dataset thus consists of 121 images from 12 wells at each of 91 time points for the 6 clones 103, 104, 109, 110, 116, 117 where blue indicates class 0 and red a class 1, respectively.

As our dataset consists of six fully independent samples, i.e. different cell clones, we split the data into training, validation and test set with a ratio of 3:2:1. We performed the splitting six times, such that each clone is used as a test set once. For every split, each clone only appears exclusively in either training, validation, or test set and never in more than one. See Table ??. This prevents unintended information leakage due to the memorization of cells from a clone across the splits. Given that the sample size could not be easily increased due to cost and scarcity of material, we performed the maximally meaningful data splitting to ensure the best possible generalizability.

| Split | Train | Validation | Test |
| --- | --- | --- | --- |
| 1 | 104, 109, 110 | 116, 117 | 103 |
| 2 | 103, 109, 117 | 110, 116 | 104 |
| 3 | 103, 110, 116 | 104, 117 | 109 |
| 4 | 104, 116, 117 | 103, 109 | 110 |
| 5 | 103, 110, 117 | 104, 109 | 116 |
| 6 | 104, 109, 116 | 103, 110 | 117 |

### 2.2. Network training

After initial experiments with different models, the results of which can be seen in Table ??, we finetuned an EfficientNet-V2-S^20^ on the training subset, pretrained on Image-Net from torchvision 0.20.1+cu124 with PyTorch version 2.5.1+cu124. Preliminary runs showed better results for EfficientNet-V2-S than for the other tested models, see Table ??. Computational costs of all utilized models are shown in Table ??. It was trained for 15 epochs with a batch size of 64, cross entropy loss and SGD with momentum of 0.9 and learning rates between 0.01 and 0.0001. Random flipping and random cropping to 420 pixels were applied to training images. Training images for the initial set of experiments were augmented with brightness (in [0.9, 1.1]) and contrast (in [0.9, 1.1]) variations. Cropped input images will be referred to as patches in the following. Training was done independently for each of the used time points and repeated for all six data splits in Table ?? with five different seeds, resulting in 30 pairs of split/seed. A model was selected on the validation set of each split using highest patch accuracy per epoch over 15 epochs, with one final model for each of the 30 split/seed-pairs. These were then evaluated for accuracy on the test set of each split.

| Model | patch-level accuracy at 72h | clone-level accuracy at 72h |
| --- | --- | --- |
| Resnet-50 | $69.6 \pm 34.5$ | 70.0% |
| Wide-Resnet-101 | $71.0 \pm 29.7$ | 80.0% |
| Swin-V2-S | $51.9 \pm 35.8$ | 60.0% |
| ViT-B-32 224 | $56.4 \pm 32.5$ | 60.0% |
| LogReg SGD v1 | $49.5 \pm 22.3$ | 43.3% |
| LogReg SGD v2 | $48.1 \pm 14.9$ | 40.0% |
| EfficientNet-V2-S | $77.0 \pm 29.3$ | 83.3% |

| Models | GFLOPS |
| --- | --- |
| EfficientNet-V2-S | 8.37 |
| ResNet-50 | 4.09 |
| WideResNet-101 | 22.75 |
| SwinTransformer-V2-S | 11.55 |

### 2.3. Methods for the analysis of the trained model

Layer-wise Relevance Propagation (LRP) We applied LRP to the convolution layers once using LRP-β with *β* = 0,^21^ and once, using LRP-adaptive-β with *β*_*max*_ = 2.^?^ The last linear layer employed LRP-ε with *ϵ* = 10^−3^. Bias terms were ignored in the backward pass computation of the network as common for LRP modified gradient approaches. LRP was chosen due to its higher faithfulness compared to smoothed gradient methods.^?^

Structure factor The structure factor *S(q)*, where q = 1*/r* and r is the radial distance,,^22^ was calculated at certain time points for all images *I(x, y), x, y ∈ {1* … *1024}* in the following manner:

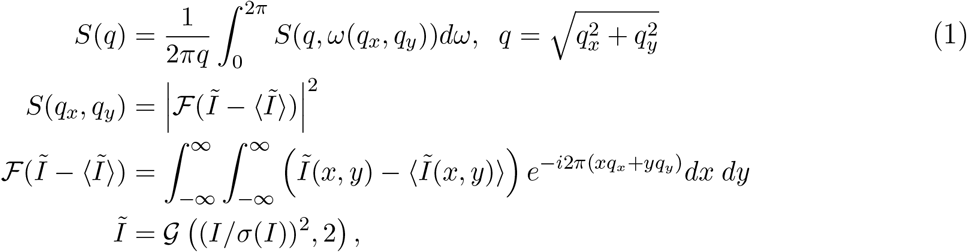

where 𝒢*(I, 2)* is a 2D Gaussian blur with standard deviation 2 pixels, ⟨ *I* ⟩ is the mean of *I* σ(*I*) is the standard deviation of *I*. The 2D Fourier transform *F(I)* has been calculated with FFT using the Matlab function fft2.

The structure factor is a measure of the spatial correlations in live/dead cell matter distribution. Figure 5 shows the results.

## 3. Results

We show at first the predictive performance measurement. Afterwards we analyze the features which the model uses to arrive at a prediction.

### 3.1. Predictive Performance

In this study, we aimed to classify six patient-derived clones of induced pluripotent stem cells (iPSC) according to their respective level of CXCR4 expression on day 4 of differentiation. This gene expression level was used as a proxy for successful differentiation towards functional β-cells. Figure 2 shows representative crops of phase-contrast images of a clone of each class at different time points during early differentiation as used in this study. Representative images for all six clones can be found in supplementary material^a^. iPSC clones were divided into two groups according to their CXCR4 expression levels with threshold of 84.0%. This threshold is based on the results of a gene expression analysis of the 6 clones, as described in section 2.1.

**Fig. 1.**
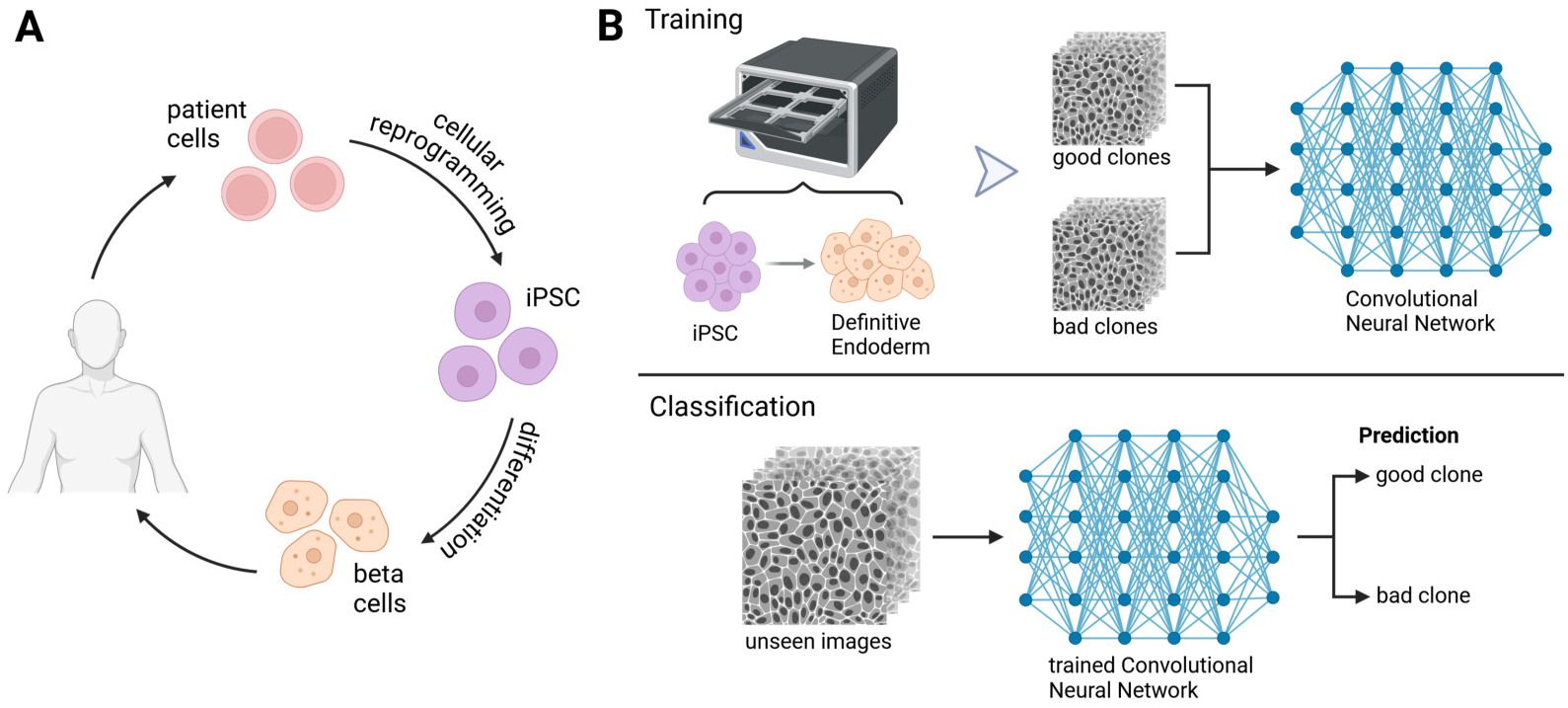
(A) Cell therapy overview for T1D. Adult cells are taken from a patient and pluripotency is induced by cellular reprogramming in vitro, resulting in patient-specific iPSC. These iPSC are then differentiated into functional β-cells before they are transplanted into the original patient again. (B) Graphical abstract showing Deep Learning approach to classify phase-contrast images from different iPSC clones. First, images are taken with IncuCyte until cells reach Definitive Endoderm (DE) stage. These images are used to train a deep neural network. Later, images of an unseen clone are fed into the trained network to get a prediction of either good (class 1) or bad (class 0) clone. Figure created in BioRender.

**Fig. 2.**
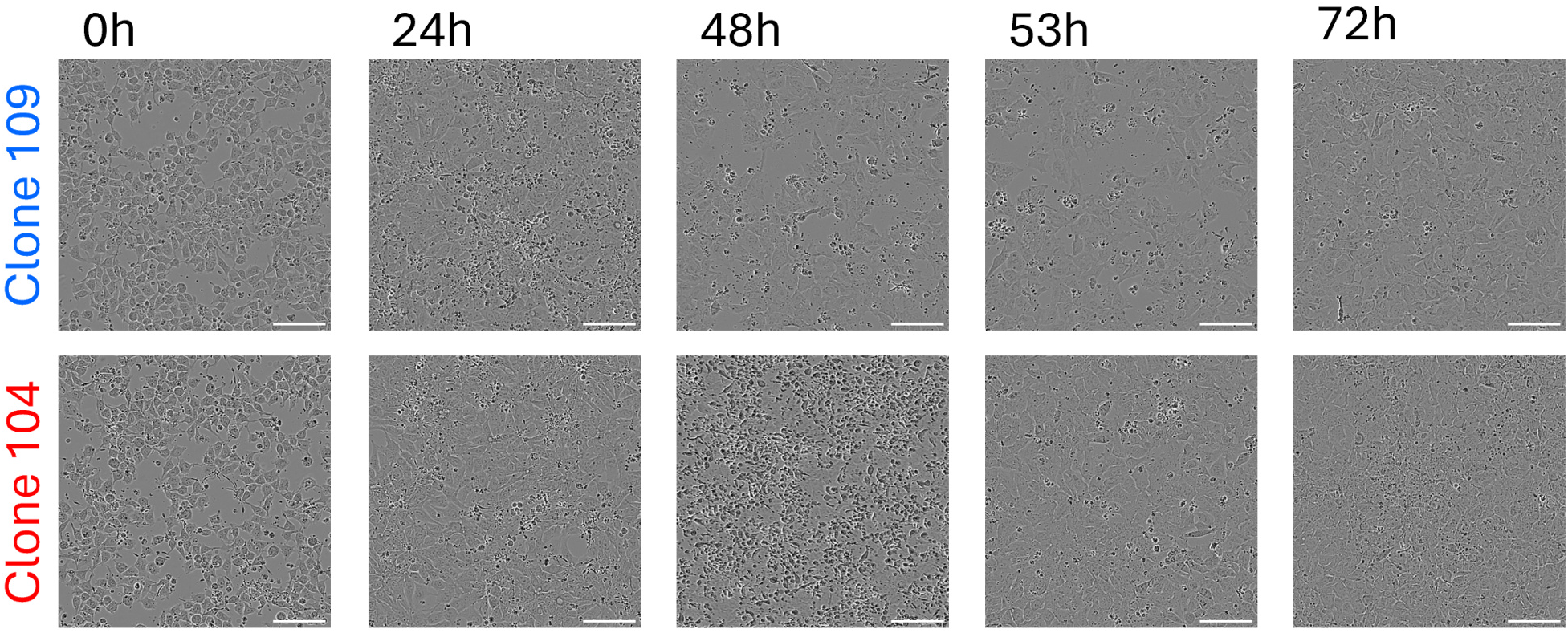
Representative sample image crops of a class 0 iPSC clone (clone 109) and a class 1 iPSC clone (clone 104). Scale bars, visible when zooming in, represent 100 *µ*m. An enlarged version of the figure can be found in the supplementary material.

**Fig. 3.**
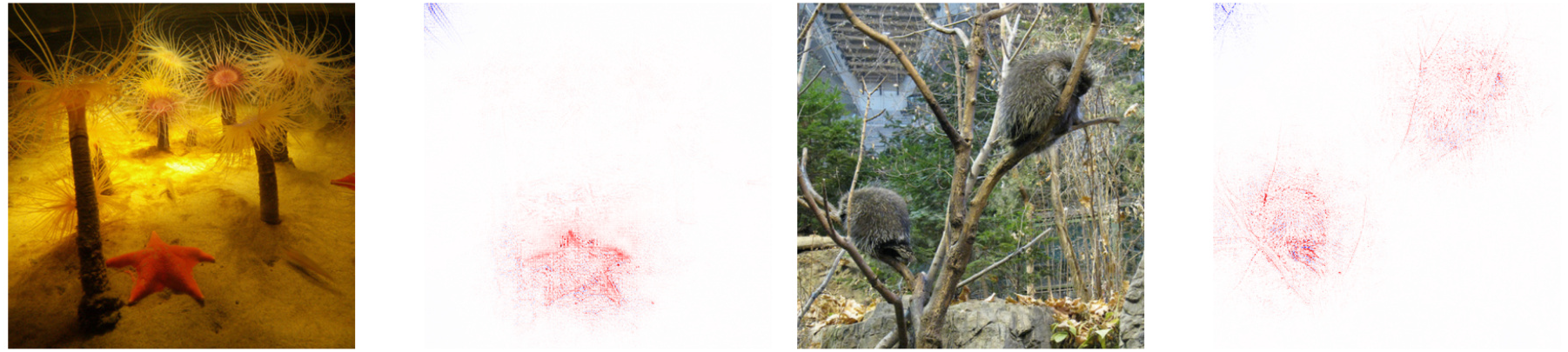
LRP heatmaps for EfficientNet-V2-S for a network pretrained on ImageNet, using images from its validation set, highlighting relevant predictive areas of the image. This shows the LRP-adaptive-β variant. Predicted classes are: starfish, porcupine. All images are from the Imagenet dataset.^23^

We report two metrics evaluating accuracy on the test set of each split, the patch accuracy and the clone accuracy. Next we describe how these are computed. At first, we draw *m* recorded images from each clone at random positions. Each image is further cropped down due to their high resolution to obtain a patch. Each patch is passed through the model with n different split-seed pairs to give m × n individual patch results p_ij_ ∈ {0, 1}, where *i* ∈ {1 … *m*} and *j* ∈ {1 … *n*}. For each split-seed pair, we calculate the seed result 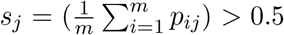 and summing over the seed results we obtain the clone accuracy 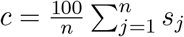 We also calculate the patch accuracy 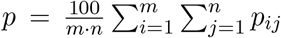 Specifically, patch accuracy is calculated as the percentage of correct predictions over all m × n patch/seed-pairs. Clone accuracy in contrast, is the percentage of correct classifications of a single clone over 5 seeds as obtained by majority vote. The random guess level for both patch and clone accuracy is 50%. This is given for clone accuracy due to an equal number of positive and negative labeled clones. For patch accuracy this was ensured by drawing an equal number of patches from each clone. Table 5 shows the results.

| measure/time | 24h | 48h | 53h* | 72h | 76h* |
| --- | --- | --- | --- | --- | --- |
| patch | 74.3 | 74.3 | 81.0 | 77.0 | 77.3 |
| accuracy (%) | $\pm 38.9$ | $\pm 39.1$ | $\pm 18.6$ | $\pm 29.3$ | $\pm 33.1$ |
| clone accuracy (%) | 76.7 | 73.3 | 96.7 | 83.3 | 76.7 |

Prediction at iPSC stage (0 hours) was not possible and resulted in random performance. This negative result is biologically plausible, as despite having several differentially expressed genes already before initiation of differentiation, visible features, such as morphology, of all iPSC clones are still too similar. The fluctuation in test accuracy does not show a clear trend over time and is likely an effect of our low effective sample size of six.

### 3.2 Insights into the cues the model is using for predictions

Initial attribution map inspection We made two observations from LRP attribution maps. Examples using LRP-β are shown in Figure 4.

**Fig. 4.**
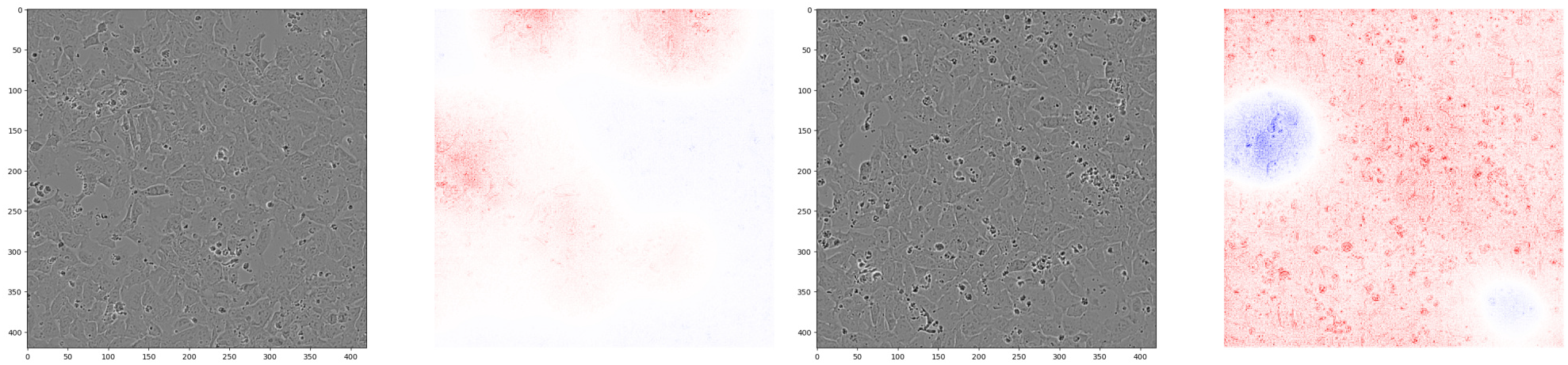
Image and LRP heatmaps for the initial model at 53 hours. Left: predicted as negative (0), thus hole-type evidence (see observation (O2)) for class 0 is in red. Right: positive prediction as class 1, thus hole-type evidence for class 0 is in blue. Evidence for the positive class covers wide areas (O1). See supplementaly material for more figures.

**Fig. 5.**
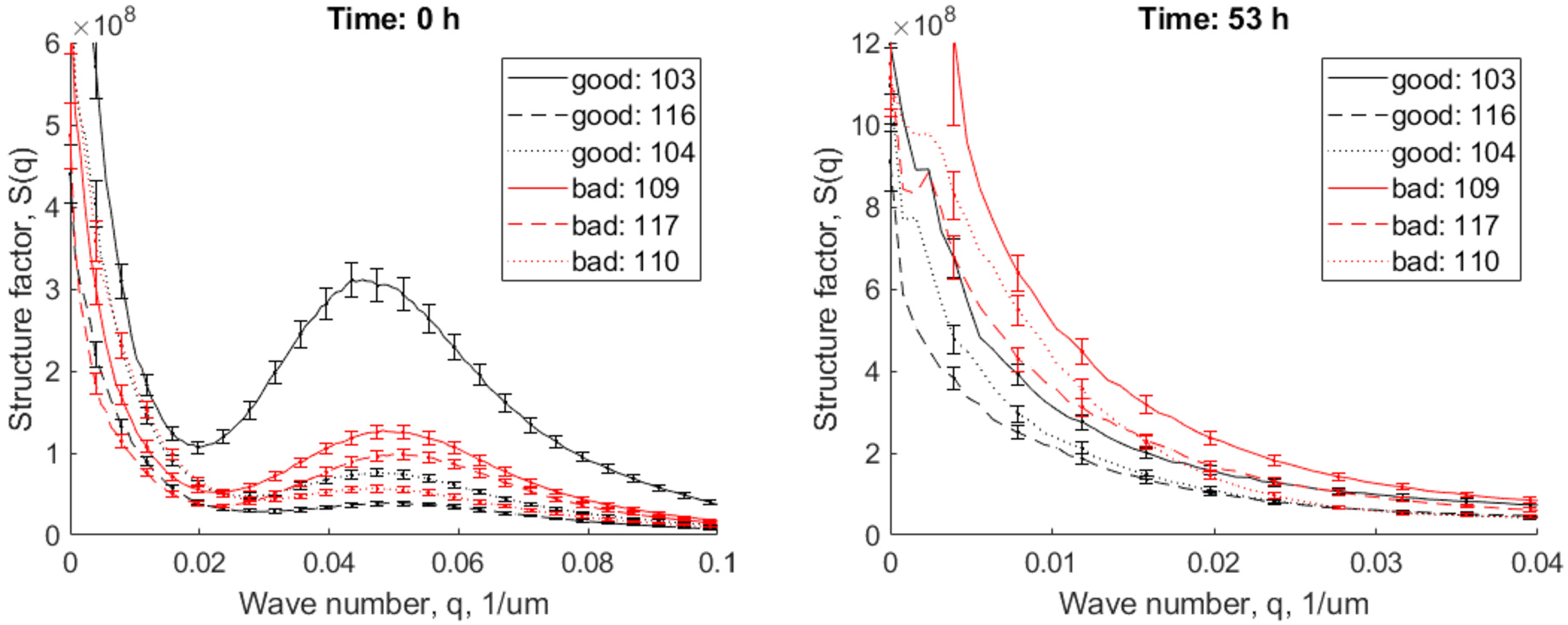
Structure factor analysis of all images at time 0 hours and 53 hours, respectively. At time 0 there is no distinct difference between the structure factor of class 0 (red curves) and class 1 (black curves) images. There is a striking peak for all clones around wave number 0.05, which is approximately 20 *µ*m. This corresponds to the size of a single cell in these images. At time 53 hours, the peak at wave number 0.05 is gone, but there is a distinguishable difference between class 0 and class 1 images at wave numbers between 0.005 and 0.02, corresponding to radial distances of 50 - 200 µm.

(O1) When predicted as positive, evidence for the positive class covers larger areas, compared to attribution maps computed for Image-Net trained models.

(O2) Evidence for the negative class 0 are clean holes without cells. Clean refers to low texture content.

Regarding (O1) we note that our dataset typically shows a dense matrix of relevant cells covering most of the image, unlike natural images from ImageNet which show objects of varying sizes, and, also unlike a number of samples of HE-stained cancer histopathology images which in many regions contain Haematoxylin-stained nuclei sparsely spread within large areas of Eosin-stained extracellular material. ?? shows figures for the original ImageNet-trained network for comparison of the areas covered by evidence. This shows the explanations produced by LRP on samples from ImageNet for which the class-relevant cues are well understandable for non-experts.

Regarding (O2) the model seems to learn the absence of a cue. Hole-type structures with a larger area showing absence of cells in it indicates lower cell confluency, which is an indication of lower quality for a cell type meant to form a closed monolayer.

Structure factor analysis To understand the typical scales of structure, we calculated the structure factor S(q) for all images of a time point after correcting for brightness variance. Figure 5 shows the results of the Fourier analysis at times 0 hours and 53 hours. At 53 hours, there is a region in the graph where there is a visible difference between the good and bad clones’ structure factor. The structure factor *S*(*q*) measures periodic correlations over different length scales r = 1/q. A larger value of the structure factor in a range of wave vectors *q* signifies that in the corresponding range of length scales, there is a stronger correlation of intensities. Thus, the measurement that the bad clones have a significantly higher S(q) in the range *r* ∈ [50, 200] *µ*m signifies that there are more persistent regions of high or low intensity over the range of 50 to 200 *µ*m. Interestingly, the LRP heatmaps show red and blue regions over similar length scales, meaning our model likely learned these differences between the clones over this particular scale. Looking at the results at 0 hours, there is no difference between the two groups of clones visible, but there is a very clear spike at a scale of approximately 20 µm, the size of a single cell in our images.

Search for global cues Having seen that the cells cover large areas, we are inspecting global statistics and their relation to cues for the model.

Obvious candidates are the mean and variance per patch. Figure 6 shows the impact of cell washing on these variance. One can see that it reduces the variance notably in 2 of the 3 washing times, and still visibly in the last washing after 3d04h00m. Note that the images were taken inside a dark chamber, so there is no influence from stray light, and the changes reflect dynamics of cells and the impact of washing. One notable consequence of cell washing is the removal of darker cell debris.

**Fig. 6.**
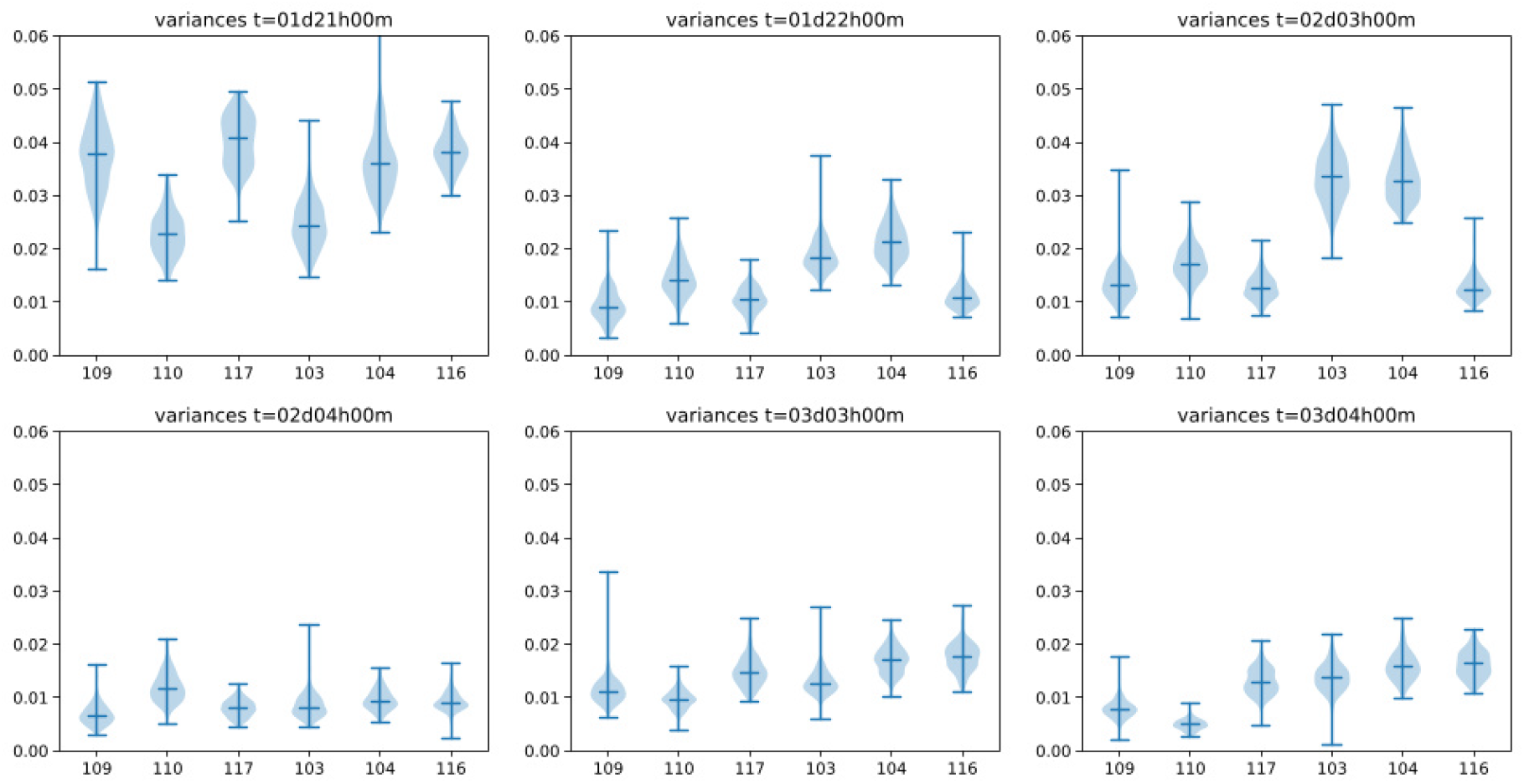
This shows 3 pairs of variances at 3 pairs of times before and after cell washing. Means are shown in the supplementary material.One can see an impact of the procedure.

In the next step we measured the extent of correlation of the attribution maps to the patch mean, the patch variance, and pixel-wise edge strength. We did this by measuring Spearman rank correlation to the pixel-wise intensity, which sums up to the mean, to the pixel-wise square deviation from the mean, which sums up to the patch variance, and to pixel-wise edge norms obtained from a Prewitt filter^24^ applied to the image patches. We obtain one Spearman rank correlation per image, and report the mean over 1200 images of the test set for each of 3 correlations. Table 6 shows the results. We see that pixel-wise variance is notably rank correlated to the attribution maps.

| Attribution map | Patch mean | Patch variance | Edge norm |
| --- | --- | --- | --- |
| LRP-adaptive-β | $0.03 \pm 0.02$ | $0.35 \pm 0.15$ | $-0.01 \pm 0.05$ |
| LRP-β | $0.01 \pm 0.01$ | $0.44 \pm 0.19$ | $0.01 \pm 0.05$ |

Variance normalized prediction We can see in Figure 6 that the variance is in parts different between the positive and negative clones. It is open whether the patch variance is causally related to the labels. To investigate, we devised an alternative protocol, in which we removed the brightness and contrast augmentation for training and used for all data subsets a patch-wise normalization of the standard deviation. The supplementary material shows more LRP attribution map for this variant at 53 hours, confirming observations (O1) and (O2). Table 7 shows partial improvements in patch accuracy and clone accuracy compared to the results in Section 3.1, namely for data sets from those times after washing which have less debris.

| measure/time | 24h | 48h | 53h* | 72h | 76h* |
| --- | --- | --- | --- | --- | --- |
| patch | 70.9 | 71.0 | 90.3 | 76.0 | 83.2 |
| accuracy (%) | $\pm 40.4$ | $\pm 37.6$ | $\pm 8.6$ | $\pm 34.0$ | $\pm 29.9$ |
| clone accuracy (%) | 73.3 | 70.0 | 100.0 | 83.3 | 80.0 |

This initial result suggests as tradeoff that brightness augmentation is better for images with more debris, while variance normalization performs better when the amount of debris is reduced by washing.

## 4. Discussion

In this study, we present encouraging results for accurate prediction of stem cell differentiation outcomes based on phase-contrast images. Similar studies exist, but are focusing on different cell fate decisions.^17,18,25^

Our model can classify high- and low-performing iPSC clones already at 24 hours after induction of the differentiation process, which makes it highly relevant for early selection of cell clones. In further distinction to other studies we show, that our model decision is based on biologically relevant image features, such as spatial distribution of the cells as well as brightness variance of the image using explainable-AI and traditional image analysis methods.

Simplicity in the imaging modality is both the key advantage and key challenge of our approach. On the positive side, a simple imaging technique lowers the effort and cost of analysis of cell clones when using our final model. However, phase-contrast images at 10x magnification only hold limited information about the imaged cells, when compared to staining for specific expressed proteins, and therefore make it harder for the algorithm to find meaningful features in them.

A major limitation of this study is the low effective sample size of six, which is a common problem in biomedical settings due to resource limitations. Even though we tried to account for this by using cross-validation and patch-wise accuracy measurements, we have to be careful when interpreting our results.^26^ Further experiments with a larger and potentially more diverse, e.g. iPSC derived from several different patients, should be done to investigate the generalizability of our results.

Overall, our results show the potential of deep learning models in combination with automated, low-effort imaging techniques as a way bridge the gap between human expertise and computational pattern recognition. In the future this could help bring cell therapy one step closer to the clinic.

## Acknowledgments

Ethical statement Induced pluripotent stem cells (iPSCs) were derived after obtaining written informed consent from the donor, in accordance with the Declaration of Helsinki and following approval by the appropriate institutional review board (IRB). All procedures were conducted in compliance with relevant national and international guidelines and regulations governing human biological materials.

Declaration of interest The authors declare that there are no conflicts of interest associated with this publication. All authors have no financial or personal relationships that could impact the findings presented.

Funding sources This project has received funding from the European Union’s Horizon 2020 research and innovation program under the Marie Skłodowska-Curie grant agreement Nº 945371.

CRediT authorship contribution statement F.S. Conceptualization, Methodology, Data curation, Software, Writing-Original draft preparation. A.B. Conceptualization, Methodology, Software, Writing-Reviewing and Editing, Supervision. V.Z. Investigation. V.S. Conceptualization, Resources, Supervision. H.S. Conceptualization, Supervision. A.M-S. Conceptualization, Funding acquisition, Supervision. D.K.D. Conceptualization, Formal Analysis, Writing-Reviewing and Editing, Resources, Supervision.

## Appendix

### 4.1. Figure 2 of main article enlarged

**Fig. 7.**
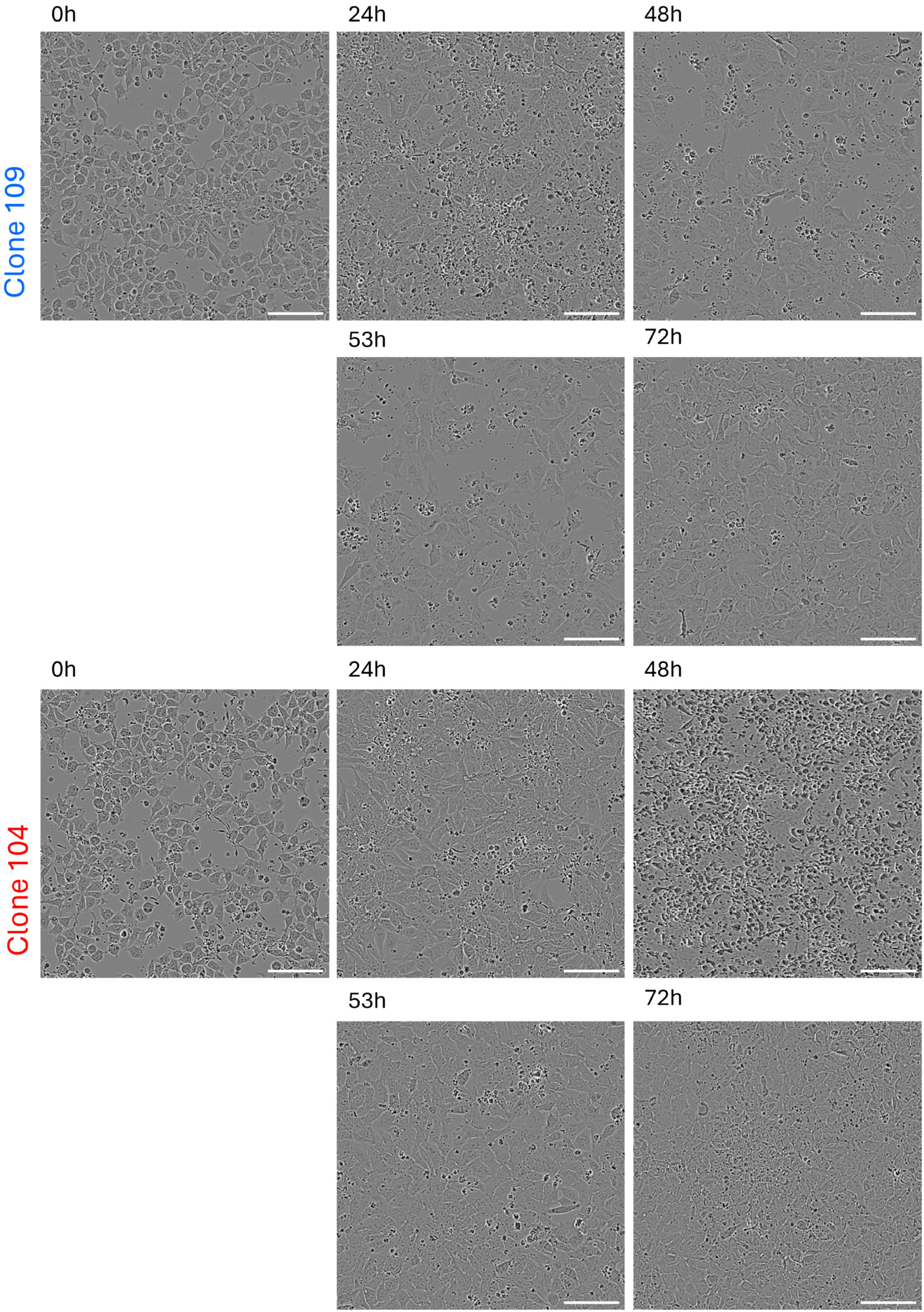
Representative sample image crops of a class 0 iPSC clone (clone 109) and a class 1 iPSC clone (clone 104). Scale bars represent 100 µm.

### 4.2. Sample images of all iPSC clones

**Fig. 8.**
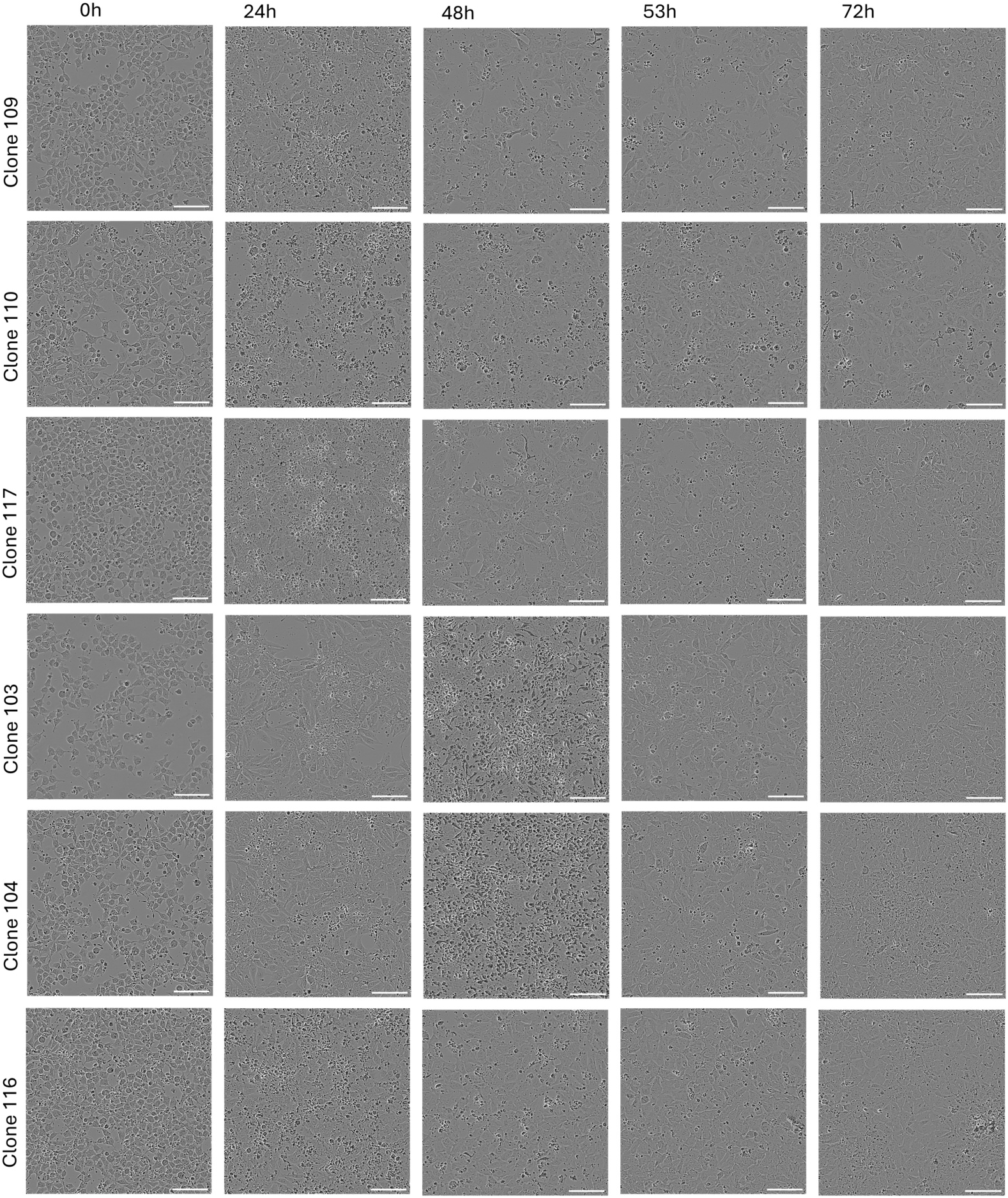
Representative sample image crops of all six imaged iPSC clones over different time points. The upper three rows show clones below the threshold, class 0, the lower rows display clones with high CXCR4 expression, class 1. Scale bars represent 100 µm.

### 4.3. More LRP heatmaps for the initial, brightness-augmented model at 53 hours for LRP-adaptive-beta

**Fig. 9.**
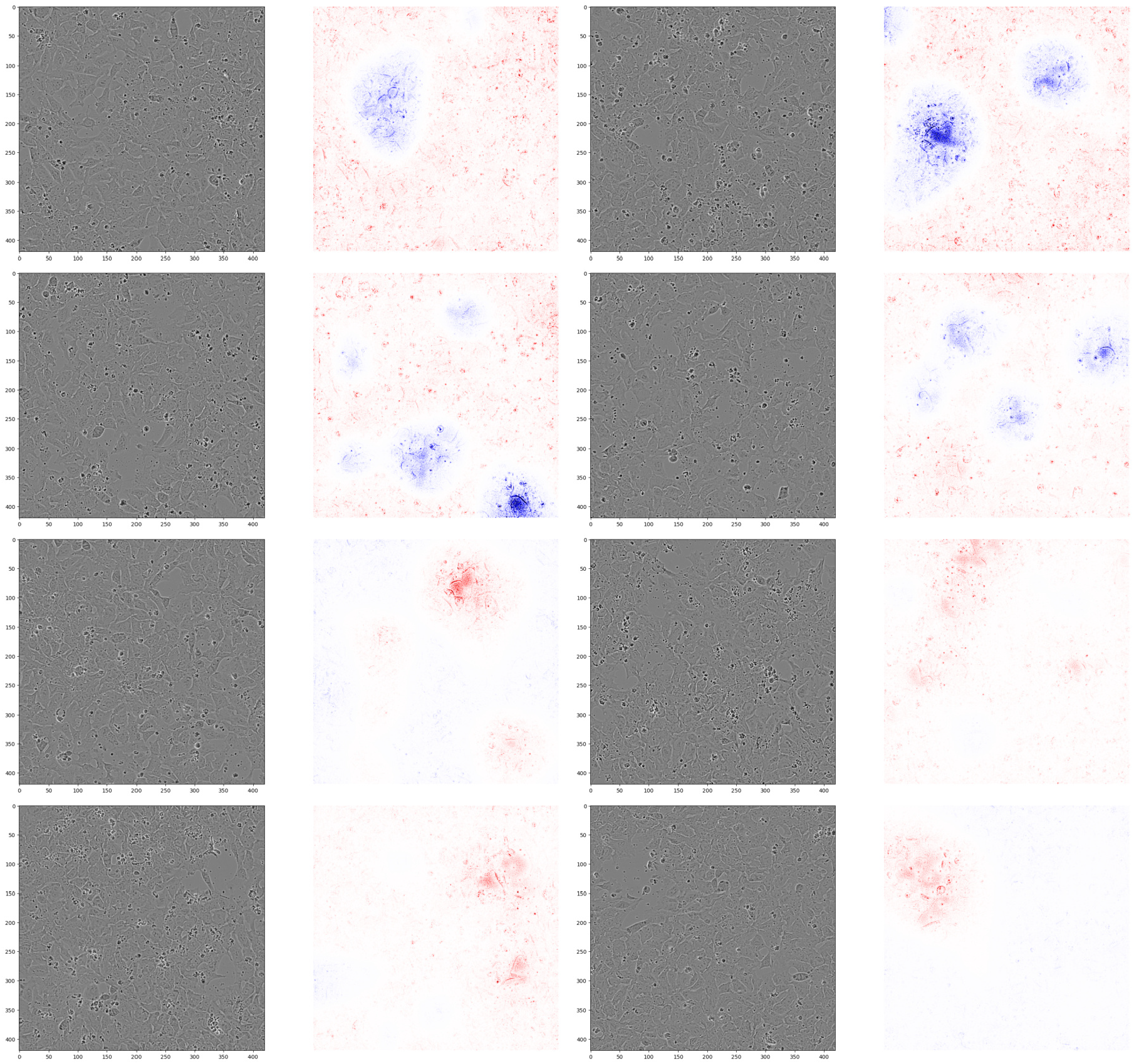
Additional images and LRP heatmaps for the initial, augmented model using LRP-adaptive-β. First and second row: predictions for class 1. For these, evidence for class 0 is in blue. Third and fourth row: predictions for class 0. For these, evidence for class 0 is in red. Time is 53 hours.

### 4.4. LRP heatmaps for the initial, brightness-augmented model at 53 hours for LRP-(nonadaptive)-beta

**Fig. 10.**
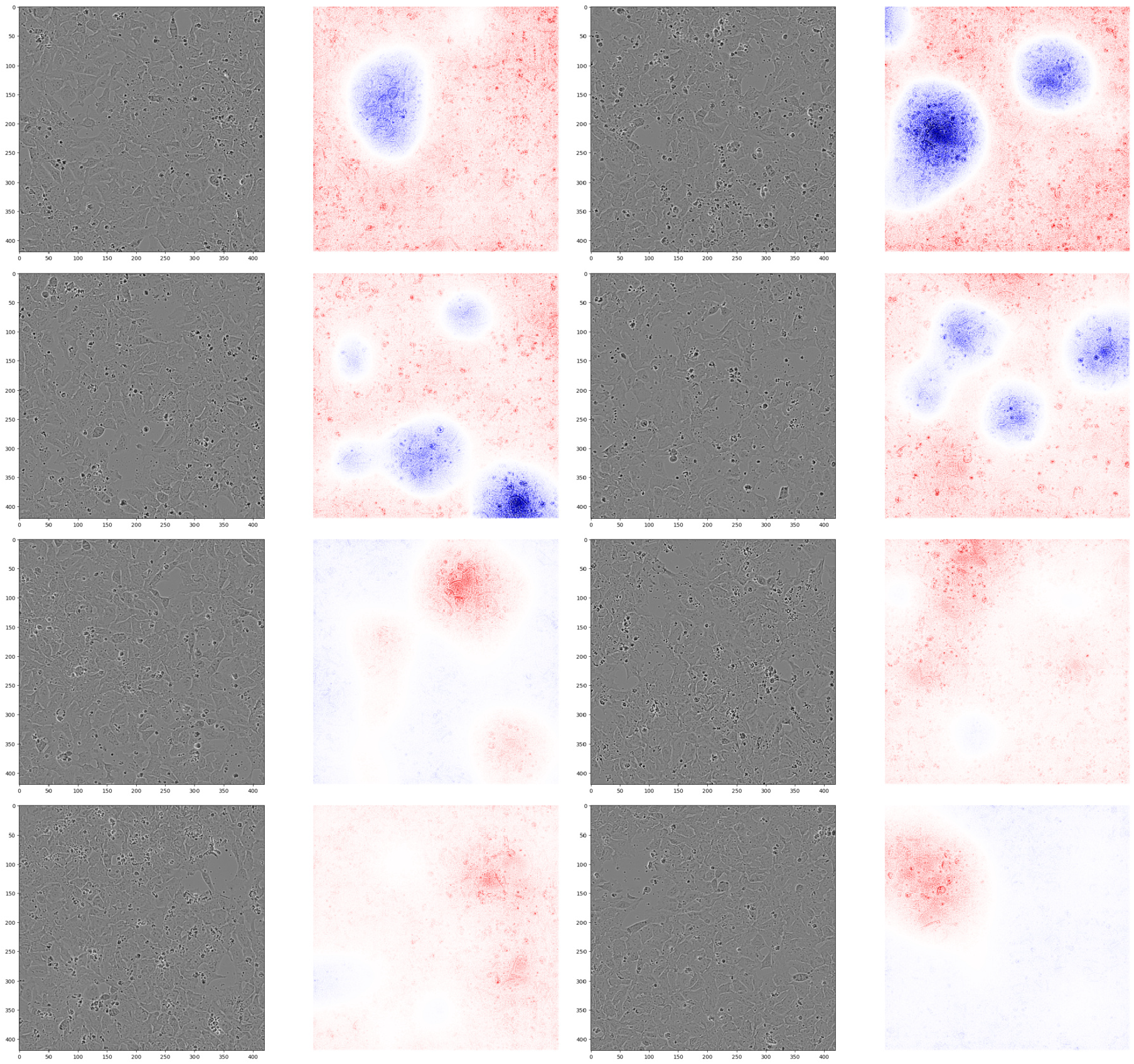
Additional images and LRP heatmaps for the initial, augmented model using LRP-β, *β* = 0. First and second row: predictions for class 1. For these, evidence for class 0 is in blue. Third and fourth row: predictions for class 0. For these, evidence for class 0 is in red. Time is 53 hours. The images are the same as those in Appendix 4.3.

### 4.5. Means before and after 3 cell washing times

**Fig. 11.**
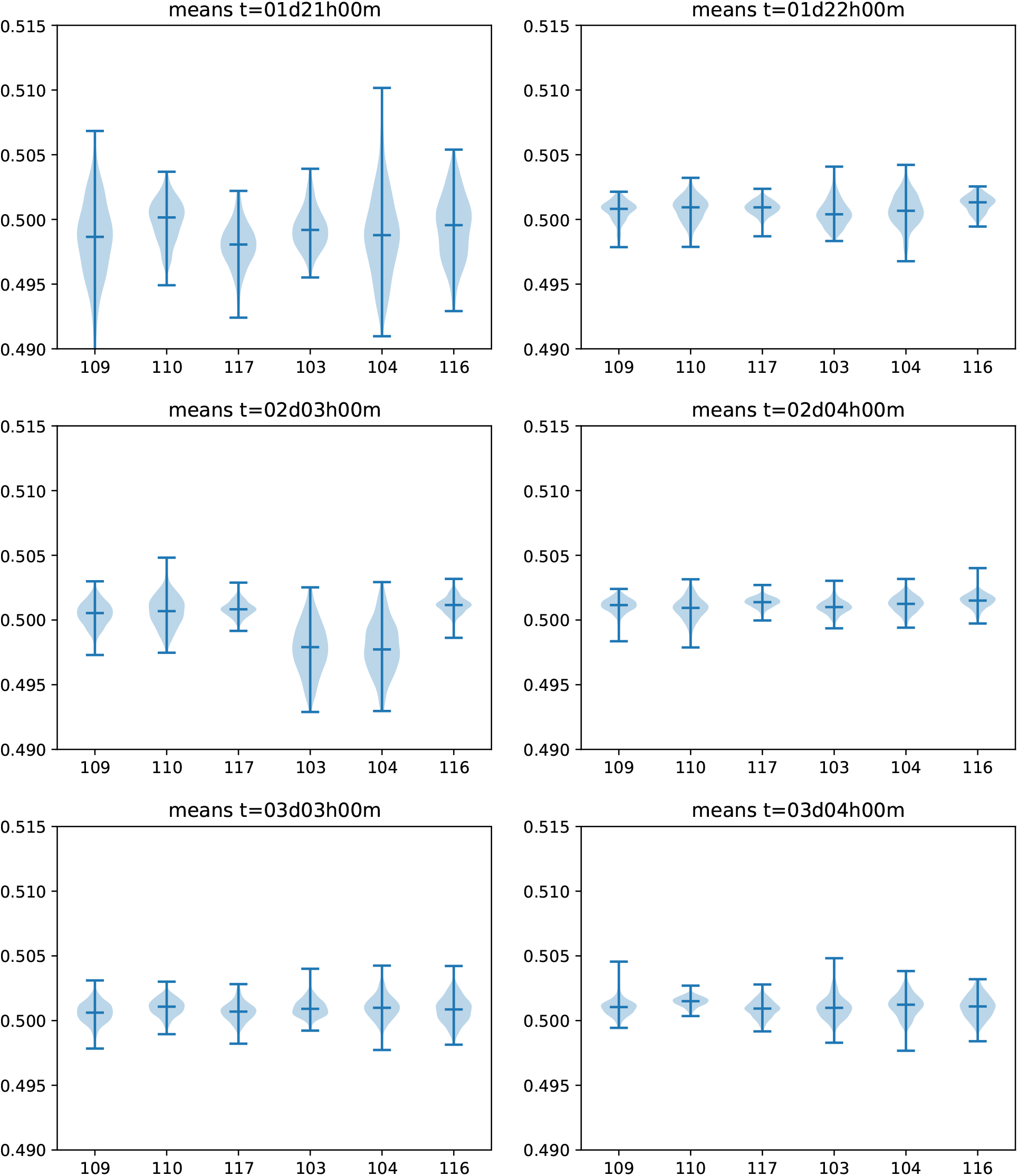
This shows 3 pairs of means at 3 pairs of times before and after cell washing. The time pairs are 01d21h00m/01d22h00m, 02d03h00m/02d04h00m and 03d03h00m/03d04h00m.

### 4.6. Mean and Variance in 12 hour intervals

**Fig. 12.**
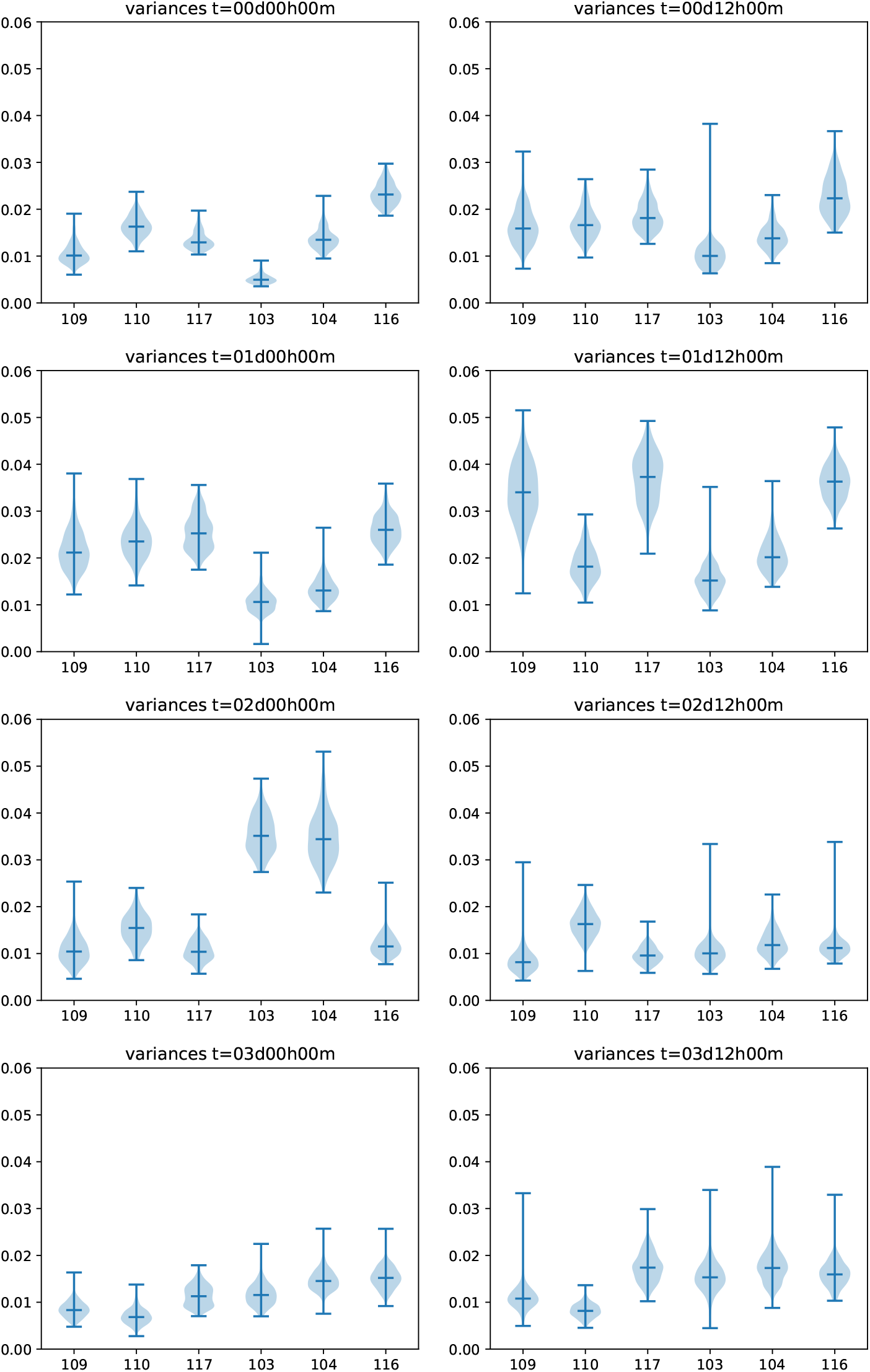
This shows variances in intervals of 12 hours.

**Fig. 13.**
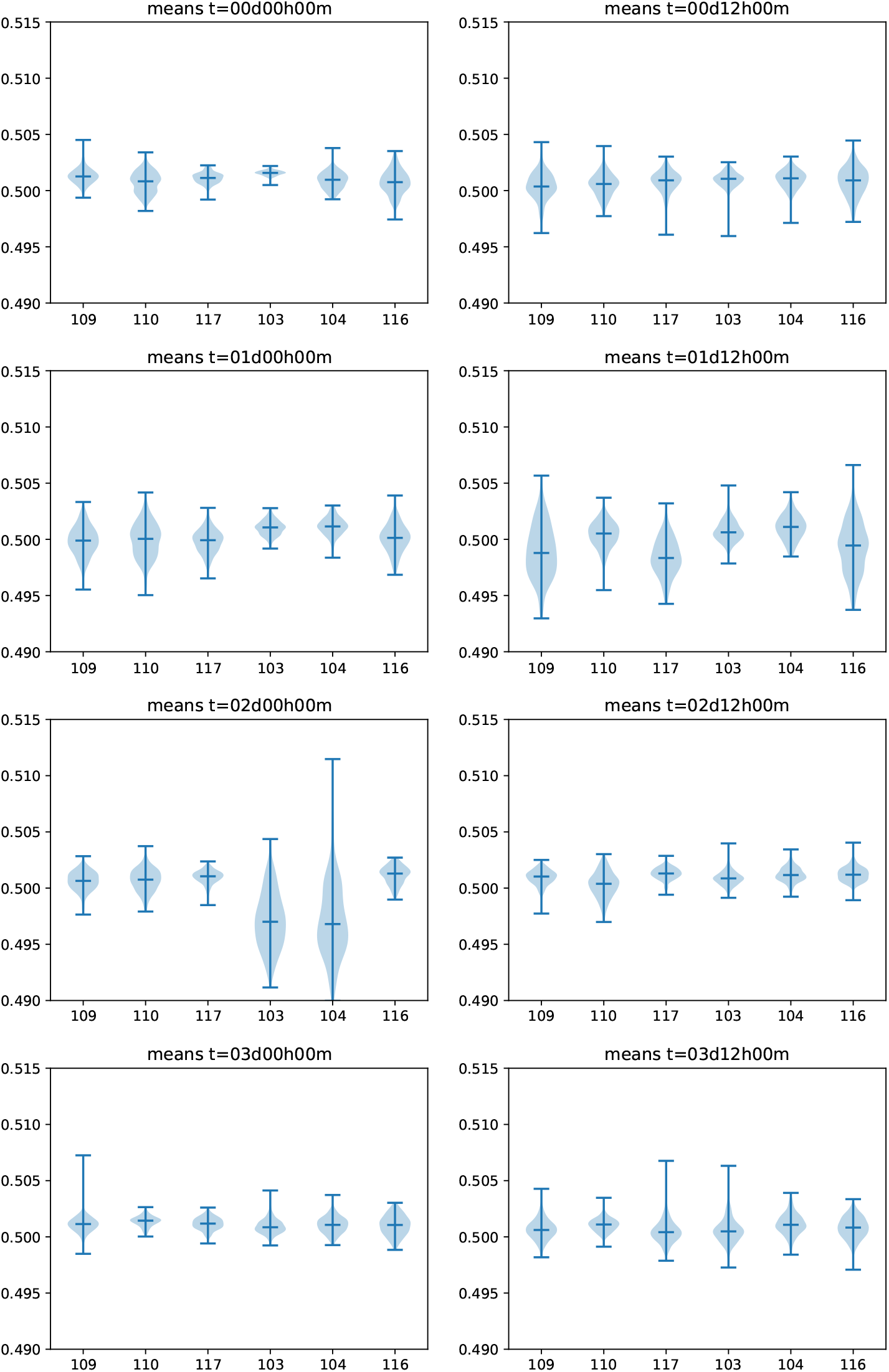
This shows means in intervals of 12 hours.

### 4.7. LRP heatmaps for the variance-normalized model at 53 hours

**Fig. 14.**
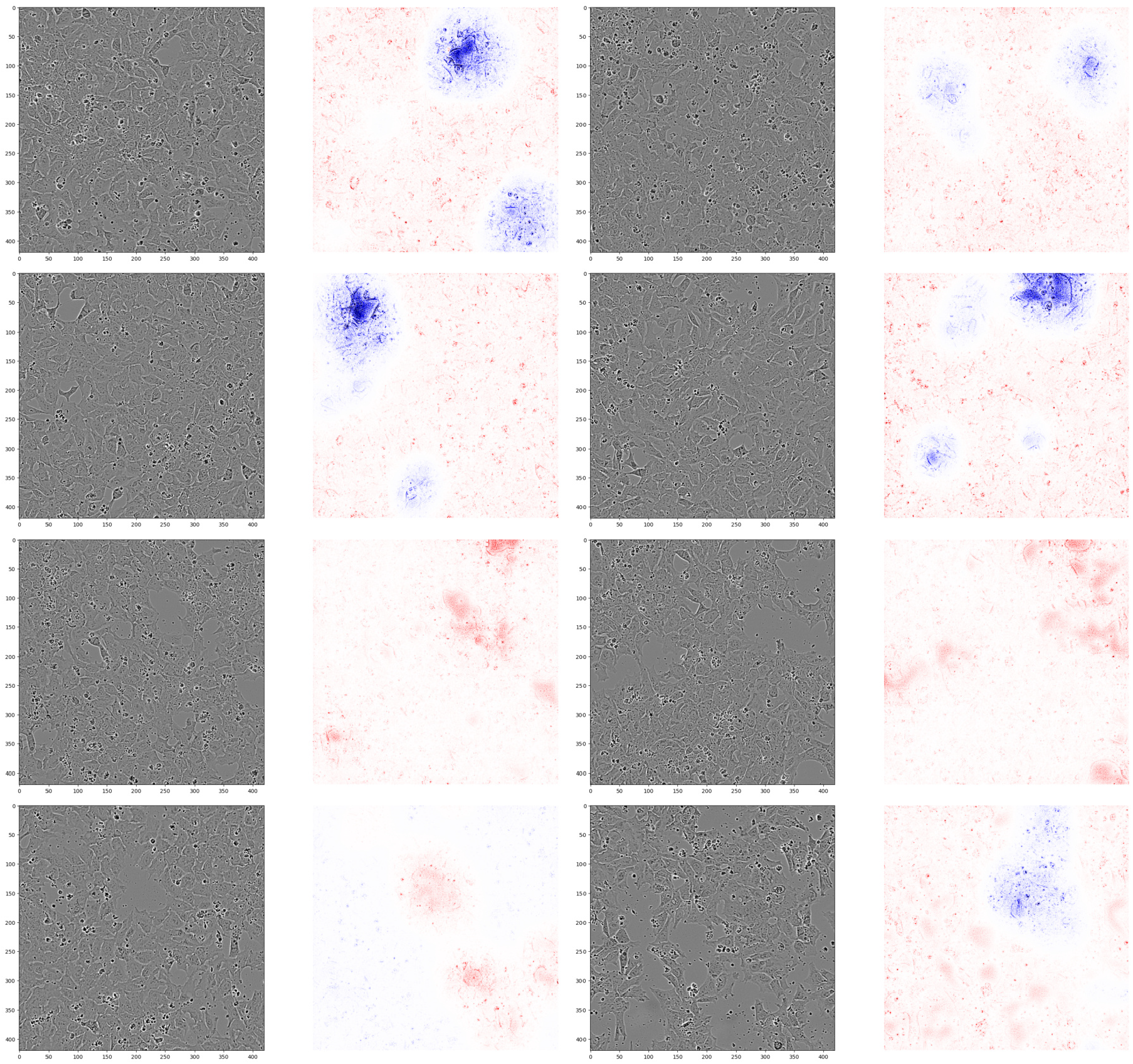
Images and LRP heatmaps for the variance-normalized model using LRP-adaptive-β. Time is 53 hours. First and second row: predictions for class 1. For these, evidence for class 0 is in blue. Third and fourth row: predictions for class 0. For these, evidence for class 0 is in red. One can see again as evidence for the class 0 holes between cells. The images are different from those in Appendix 4.3, because forward-pass predictions differ between the two models, and thus it makes sense to select different images for showing exemplary cases.

### 4.8. Exemplary side-by-side comparison LRP and smoothed gradient-based explanations

In order to explain why we did not chose to use gradient-based explanations for our model decisions, we show here a side-side comparison to demonstrate the difference in what is highlighted. There are technical reasons to use LRP heatmaps, due to their high faithfulness to the model prediction in the sense of faithfulness measures in XAI,^27,28^ and the amount of gradient-shattering noise^29^ inherent in gradient-based methods.

We used here smooth-grad, smooth-gradient times input and vargrad, all with 75 noised copies, and a noise standard deviation of *σ* = 0.5. The number of copies 75 was chosen because it is close to the limit of the GPU memory which allowed for one image to compute within one mini-batch of copies.

**Fig. 15.**
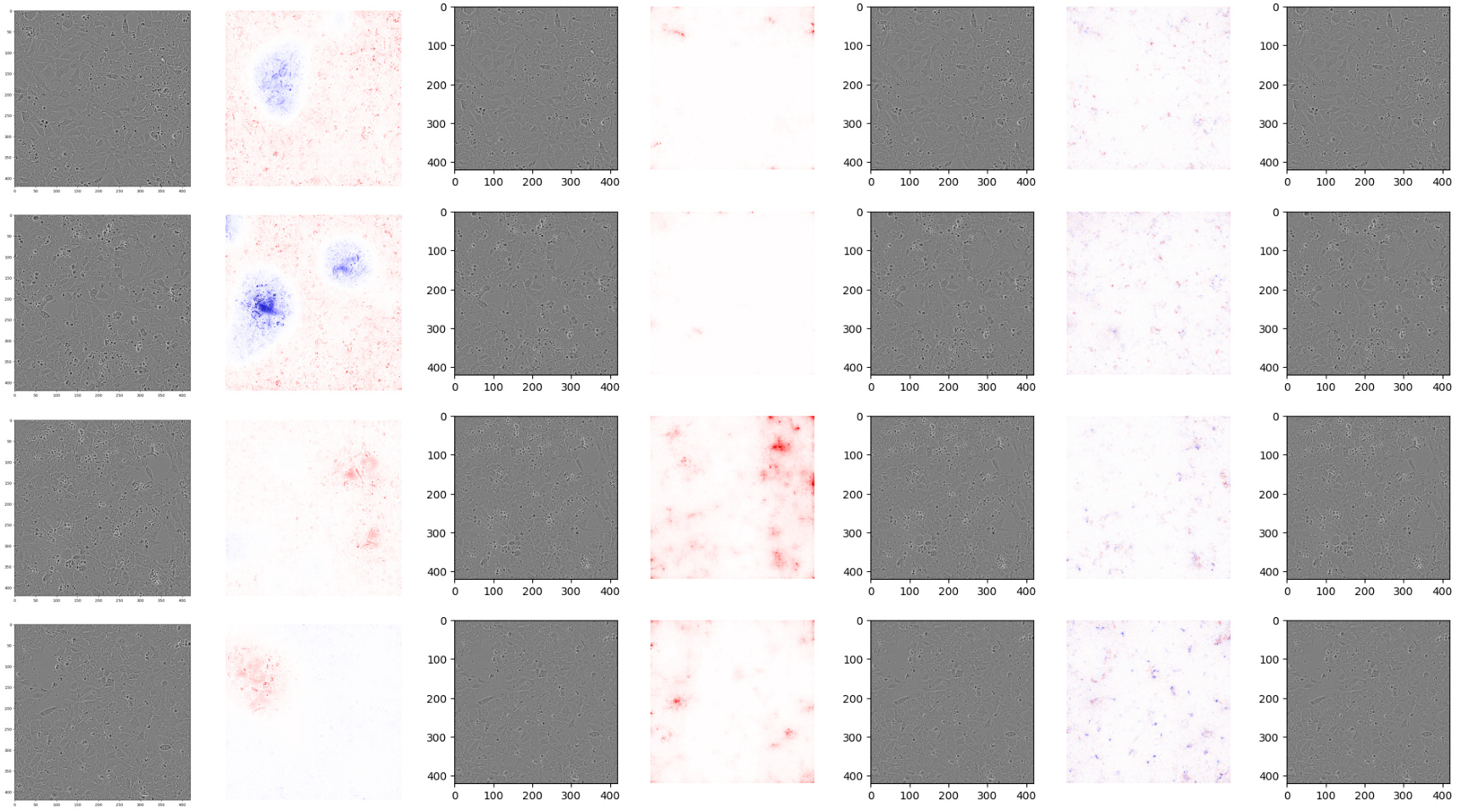
Each row shows for the same input image: LRP-Adaptive-*β*, smooth-grad, smooth-gradient times input and vargrad. Images are those from the first and last row from Appendix 4.3.

## Footnotes

a Supplementary material is available at https://doi.org/10.1101/2025.05.22.652867

## Notes

### Competing Interest Statement

The authors have declared no competing interest.

### Summary of Updates

Updated acknowledgement section; formatting updated as per target conference template

## References

1. N. C. D. R. F. Collaboration, Worldwide trends in diabetes prevalence and treatment from 1990 to 2022: a pooled analysis of 1108 population-representative studies with 141 million participants, Lancet 404, 2077 (2024).

2. J. A. Bluestone, K. Herold and G. Eisenbarth, Genetics, pathogenesis and clinical interventions in type 1 diabetes, Nature 464, 1293 (2010).

3. M. A. Atkinson, G. S. Eisenbarth and A. W. Michels, Type 1 diabetes, Lancet 383, 69 (2014).

4. V. Sordi, L. Monaco and L. Piemonti, Cell therapy for type 1 diabetes: From islet transplantation to stem cells, Horm Res Paediatr 96, 658 (2023).

5. B. Keymeulen, K. De Groot, D. Jacobs-Tulleneers-Thevissen, D. M. Thompson, M. D. Bellin, E. J. Kroon, M. Daniels, R. Wang, M. Jaiman, T. J. Kieffer, H. L. Foyt and D. Pipeleers, Encapsulated stem cell-derived beta cells exert glucose control in patients with type 1 diabetes, Nat Biotechnol 42, 1507 (2024).

6. K. Takahashi, K. Tanabe, M. Ohnuki, M. Narita, T. Ichisaka, K. Tomoda and S. Yamanaka, Induction of pluripotent stem cells from adult human fibroblasts by defined factors, Cell 131, 861 (2007).

7. Y. Shi, H. Inoue, J. C. Wu and S. Yamanaka, Induced pluripotent stem cell technology: a decade of progress, Nat Rev Drug Discov 16, 115 (2017).

8. T. Barsby, H. Ibrahim, V. Lithovius, H. Montaser, D. Balboa, E. Vahakangas, V. Chandra, J. Saarimaki-Vire and T. Otonkoski, Differentiating functional human islet-like aggregates from pluripotent stem cells, STAR Protoc 3, p. 101711 (2022).

9. V. Volpato and C. Webber, Addressing variability in ipsc-derived models of human disease: guidelines to promote reproducibility, Dis Model Mech 13 (2020).

10. J. W. Chan and A. K. K. Teo, Replicates in stem cell models-how complicated!, Stem Cells 38, 1055 (2020).

11. H. Aljuaid, N. Alturki, N. Alsubaie, L. Cavallaro and A. Liotta, Computer-aided diagnosis for breast cancer classification using deep neural networks and transfer learning, Comput Methods Programs Biomed 223, p. 106951 (2022).

12. Y. Chen, Y. Lin, X. Xu, J. Ding, C. Li, Y. Zeng, W. Liu, W. Xie and J. Huang, Classification of lungs infected covid-19 images based on inception-resnet, Comput Methods Programs Biomed 225, p. 107053 (2022).

13. S. L. Chu, K. Sudo, H. Yokota, K. Abe, Y. Nakamura and M. D. Tsai, Human induced pluripotent stem cell formation and morphology prediction during reprogramming with time-lapse bright-field microscopy images using deep learning methods, Comput Methods Programs Biomed 229, p. 107264 (2023).

14. J. Huth, M. Buchholz, J. M. Kraus, K. Molhave, C. Gradinaru, G. V. Wichert, T. M. Gress, H. Neumann and H. A. Kestler, Timelapseanalyzer: Multi-target analysis for live-cell imaging and time-lapse microscopy, Computer Methods and Programs in Biomedicine 104, 227 (2011).

15. M. ▯ Mekuc, C. Bohak, E. Bones, S. Hudoklin, R. Romih and M. Marolt, Automatic segmentation and reconstruction of intracellular compartments in volumetric electron microscopy data, Computer Methods and Programs in Biomedicine 223 (2022).

16. K. A. D’Amour, A. G. Bang, S. Eliazer, O. G. Kelly, A. D. Agulnick, N. G. Smart, M. A. Moorman, E. Kroon, M. K. Carpenter and E. E. Baetge, Production of pancreatic hormone-expressing endocrine cells from human embryonic stem cells, Nat Biotechnol 24, 1392 (2006).

17. A. Waisman, A. La Greca, A. M. Möbbs, M. A. Scarafía, N. L. Santín Velazque, G. Neiman, L. N. Moro, C. Luzzani, G. E. Sevlever, A. S. Guberman and S. G. Miriuka, Deep learning neural networks highly predict very early onset of pluripotent stem cell differentiation, Stem Cell Reports 12, 845 (2019).

18. Y. Zhu, R. Huang, Z. Wu, S. Song, L. Cheng and R. Zhu, Deep learning-based predictive identification of neural stem cell differentiation, Nature Communications 12 (2021).

19. M. Mai, S. Luo, S. Fasciano, T. E. Oluwole, J. Ortiz, Y. Pang and S. Wang, Morphology-based deep learning approach for predicting adipogenic and osteogenic differentiation of human mesenchymal stem cells (hmscs), Front Cell Dev Biol 11, p. 1329840 (2023).

20. M. Tan and Q. V. Le, Efficientnetv2: Smaller models and faster training (2021).

21. S. Bach, A. Binder, G. Montavon, F. Klauschen, K. R. Müller and W. Samek, On pixel-wise explanations for non-linear classifier decisions by layer-wise relevance propagation, PLoS ONE 10, 1 (2015).

22. D. L. Sidebottom, Fundamentals of Condensed Matter and Crystalline Physics: An Introduction for Students of Physics and Materials Science (Cambridge University Press, 2012).

23. J. Deng, W. Dong, R. Socher, L.-J. Li, K. Li and L. Fei-Fei, Imagenet: A large-scale hierarchical image database, in 2009 IEEE Conference on Computer Vision and Pattern Recognition, 2009.

24. J. M. S. Prewitt, Object enhancement and extraction, Picture Processing and. Psychopictorics (1970).

25. M. Kim, Y. Namkung, D. Hyun and S. H. Hong, Prediction of stem cell state using cell image-based deep learning, Advanced Intelligent Systems 5 (2023).

26. F. Eitel, J. P. Albrecht, M. Weygandt, F. Paul and K. Ritter, Patch individual filter layers in cnns to harness the spatial homogeneity of neuroimaging data, Sci Rep 11, p. 24447 (2021).

27. W. Samek, A. Binder, G. Montavon, S. Lapuschkin and K.-R. Müller, Evaluating the visualization of what a deep neural network has learned, IEEE Transactions on Neural Networks and Learning Systems 28, 2660 (2017).

28. A. Binder, L. Weber, S. Lapuschkin, G. Montavon, K. R. Müller and W. Samek, Shortcomings of top-down randomization-based sanity checks for evaluations of deep neural network explanations, 2023 Ieee/Cvf Conference on Computer Vision and Pattern Recognition (Cvpr), 16143 (2023).

29. D. Balduzzi, M. Frean, L. Leary, J. P. Lewis, K. W. Ma and B. McWilliams, The shattered gradients problem: If resnets are the answer, then what is the question?, in International Conference on Machine Learning (ICML),, PMLR Vol. 70 (PMLR, 2017).

